# Wide-vergence, multi-spectral adaptive optics scanning laser ophthalmoscope with diffraction-limited illumination and collection

**DOI:** 10.1101/2020.01.29.924969

**Authors:** Sanam Mozaffari, Francesco Larocca, Volker Jaedicke, Pavan Tiruveedhula, Austin Roorda

## Abstract

Visualizing and assessing the function of microscopic retinal structures in the human eye is a challenging task that has been greatly facilitated by ophthalmic adaptive optics (AO). Yet, as AO imaging systems advance in functionality by employing multiple spectral channels and larger vergence ranges, achieving optimal resolution and signal-to-noise ratios (SNR) becomes difficult and is often compromised. While current-generation AO retinal imaging systems have demonstrated excellent, near diffraction-limited imaging performance over wide vergence and spectral ranges, a full theoretical and experimental analysis of an AOSLO that includes both the light delivery and collection optics has not been done, and neither has the effects of extending wavefront correction from one wavelength to imaging performance in different spectral channels. Here, we report a methodology and system design for simultaneously achieving diffraction-limited performance in both the illumination and collection paths for a wide-vergence, multi-spectral AO scanning laser ophthalmoscope (SLO) over a 1.2 diopter vergence range while correcting the wavefront in a separate wavelength. To validate the design, an AOSLO was constructed to have three imaging channels spanning different wavelength ranges (543 ± 11 nm, 680 ± 11 nm, and 840 ± 6 nm, respectively) and one near-infrared wavefront sensing channel (940 ± 5 nm). The AOSLO optics and their alignment were determined via simulations in optical and optomechanical design software and then experimentally verified by measuring the AOSLO’s illumination and collection point spread functions (PSF) for each channel using a phase retrieval technique. The collection efficiency was then measured for each channel as a function of confocal pinhole size when imaging a model eye achieving near-theoretical performance. Imaging results from healthy human adult volunteers demonstrate the system’s ability to resolve the foveal cone mosaic in all three imaging channels despite a wide spectral separation between the wavefront sensing and imaging channels.

**OCIS codes:** (110.1080) Active or adaptive optics; (170.4460) Ophthalmic optics and devices; (170.4470) Ophthalmology

## 1. Introduction

The adaptive optics scanning laser ophthalmoscope (AOSLO) is an important imaging tool that can achieve in vivo, near diffraction-limited visualizations of microscopic structures in the retina by compensating for the monochromatic aberrations of the eye [1].

Increasingly, adaptive optics systems are employing multiple wavelength channels for a range of imaging and vision testing applications. Most systems use different wavelengths for wavefront sensing and imaging. Systems for AOSLO microperimetry and visual psychophysics employ NIR wavelengths for wavefront sensing and tracking and deliver AO-corrected flashes of visible light. Other multi-wavelength applications include fluorescence [2–4], hyperspectral imaging [5], retinal oximetry [6] and multi-modal imaging [7]. There are two important factors to consider when designing, building, and interpreting results from a high-fidelity multi-wavelength AOSLO system: the chromatic aberrations of the eye and the chromatic aberrations of the system.

### Chromatic effects of the eye

First and foremost, the chromatic dispersion of the eye needs to be considered. The chromatic difference in defocus of the eye (longitudinal chromatic aberration, or LCA) has been extensively studied and its behavior is quite predictable and similar between individuals [8], albeit with inter-individual differences depending on the eye’s specific optical parameters (e.g. corneal curvature). The transverse chromatic aberration, or TCA, of the eye has also been studied [9,10] and recently objective techniques to measure it [11,12] and correct it [13,14] have been employed. The extent to which the high order aberrations of the eye change with wavelength is less studied. As a rule, there must be differences in high-order aberrations as the ray paths that different wavelengths of light take through the optical system to reach a focus on the retina are different. But, researchers generally agree that these differences are small and often fall within the range of measurement error of the system [15–16].

In the current study, our aim was to carefully explore the implications of using NIR wavelengths for wavefront sensing and visible light for imaging. In a diffraction-limited system, the size of the point spread function is proportional to wavelength, so the best possible outcome is that the changes in high-order aberrations between wavelengths are negligible, and the benefits of reduced diffraction would yield increasingly sharp images at progressively shorter wavelengths even if wavefront sensing was performed at long (NIR) wavelengths. In the worst possible outcome, the high order aberrations would change enough to offset the benefits of reduced diffraction when imaging at wavelengths shorter than the wavefront sensing wavelength.

### Chromatic effects of the AOSLO system

To facilitate an exploration of the implications of high-order changes in chromatic aberrations in the eye for multi-wavelength AOSLO systems, it is paramount to minimize or fully characterize the chromatic effects of the AOSLO itself. The bulk of this manuscript describes this effort. Current-generation AOSLOs are typically able to achieve high resolution over wide vergence ranges (~3 diopters). Optimal performance has been achieved by designing the relay telescopes in the AOSLO with either a non-planar design using off-axis spherical mirrors [5,17], a planar design using a combination of off-axis toroidal and spherical mirrors [18], or an on-axis, lens-based design using polarized light and polarization gating to minimize back-reflections from refractive surfaces [19]. However, to our knowledge, a similar consideration of vergence effects in the collection path of an AOSLO has not yet been described, despite the use of off-axis optical elements such as plate/wedge beamsplitters and dichroic mirrors in the collection path, which induce vergence-dependent aberrations. Here, we present a multi-spectral AOSLO design with diffraction-limited performance in both the illumination and collection paths over a 1.2 diopter vergence range. The optical design utilized all commercially available off-the-shelf optics thereby improving the cost-efficiency of our system compared to prior AOSLOs using custom-sized spherical mirrors [5] or custom toroidal mirrors [18]. To facilitate the optical alignment and construction of this AOSLO design, we developed a detailed optomechanical model and constructed a laser-cut polycarbonate stencil, which indicated the placement of all optomechanics onto an optical table. After the AOSLO was constructed and aligned, we experimentally validated the system’s resolution and collection efficiency using a phase retrieval technique and a model eye setup.

Finally, we present imaging results at the fovea in two healthy adult volunteers and quantitatively compare resolution across imaging channels using a Fourier analysis. For wavefront sensing at 940 nm, the benefits of reduced diffraction extends to images taken with shorter wavelengths. In the two subjects that were imaged using optimal confocal pinholes [20], the foveal cone mosaic was well resolved at all wavelengths (840, 680 and 543 nm). The benefits of reduced diffraction with shorter wavelengths was readily visible but images at 680 nm and 543 nm were similar in quality indicating that the effects of higher order aberrations of the eye may begin to play a small role for shorter wavelengths.

## 2. Methods

### 2.1 System overview and optical design

The schematic and optical design of the multi-spectral AOSLO system is shown in Figure 1 with a specification of components in Table 1. Light from a supercontinuum light source was separated into three imaging channels spanning different wavelength ranges (543 ± 11 nm, 680 ± 11 nm, and 840 ± 6 nm, respectively) and one near-infrared wavefront sensing channel (940 ± 5 nm). All spectral channels were aligned to be collinear using pairs of mirrors for each color channel and 3 dichroic mirrors. A 10:90 (R:T) wedge plate beam splitter was placed after the dichroic mirrors to separate the illumination and collection paths and was followed by four sets of spherical mirror-based telescopes arranged in a non-planar off-axis manner that relayed an image of the system’s entrance pupil onto the resonant scanner, galvanometer scanner, deformable mirror and, finally, the pupil of the eye. The positions and angles of all optics after the beam splitter in the illumination path (see Table 2) were optimized to minimize aberrations, mainly astigmatism, using optical design software (Zemax, LLC, Kirkland, WA) and the technique described by Dubra et al. [5]. Diffraction-limited illumination spots were achieved for a 1° field of view (FOV) across all AOSLO spectral channels, as indicated by the spot diagrams and Strehl ratios at the bottom of Figure 1.

**Table 1.**
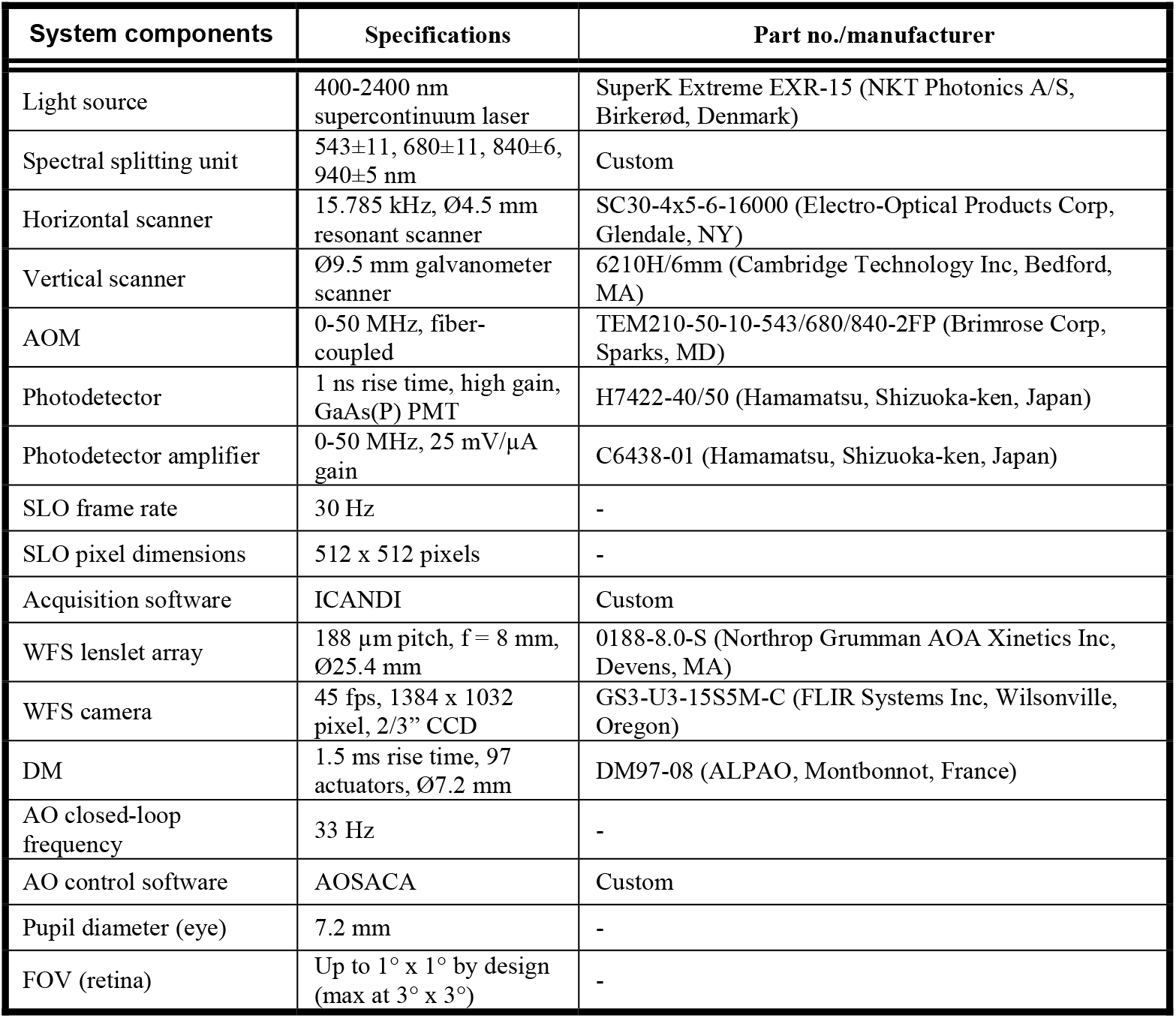
System components and hardware.

| System components | Specifications | Part no./manufacturer |
| --- | --- | --- |
| Light source | 400-2400 nm supercontinuum laser | SuperK Extreme EXR-15 (NKT Photonics A/S, Birkerød, Denmark) |
| Spectral splitting unit | 543±11, 680±11, 840±6, 940±5 nm | Custom |
| Horizontal scanner | 15.785 kHz, Ø4.5 mm resonant scanner | SC30-4x5-6-16000 (Electro-Optical Products Corp, Glendale, NY) |
| Vertical scanner | Ø9.5 mm galvanometer scanner | 6210H/6mm (Cambridge Technology Inc, Bedford, MA) |
| AOM | 0-50 MHz, fiber-coupled | TEM210-50-10-543/680/840-2FP (Brimrose Corp, Sparks, MD) |
| Photodetector | 1 ns rise time, high gain, GaAs(P) PMT | H7422-40/50 (Hamamatsu, Shizuoka-ken, Japan) |
| Photodetector amplifier | 0-50 MHz, 25 mV/μA gain | C6438-01 (Hamamatsu, Shizuoka-ken, Japan) |
| SLO frame rate | 30 Hz | - |
| SLO pixel dimensions | 512 x 512 pixels | - |
| Acquisition software | ICANDI | Custom |
| WFS lenslet array | 188 μm pitch, f = 8 mm, Ø25.4 mm | 0188-8.0-S (Northrop Grumman AOA Xinetics Inc, Devens, MA) |
| WFS camera | 45 fps, 1384 x 1032 pixel, 2/3" CCD | GS3-U3-15S5M-C (FLIR Systems Inc, Wilsonville, Oregon) |
| DM | 1.5 ms rise time, 97 actuators, Ø7.2 mm | DM97-08 (ALPAO, Montbonnot, France) |
| AO closed-loop frequency | 33 Hz | - |
| AO control software | AOSACA | Custom |
| Pupil diameter (eye) | 7.2 mm | - |
| FOV (retina) | Up to 1° x 1° by design (max at 3° x 3°) | - |

**Table 2.** Focal length, diameter, and angles of incidence on reflective optical elements of the AOSLO.

| Optical element | Focal length (mm) | Diameter (mm) | $I_x$ (deg) | $I_y$ (deg) |
| --- | --- | --- | --- | --- |
| Spherical mirror # 1 | 500 | 25.4 | 4.00 | -1.00 |
| Spherical mirror # 2 | 500 | 25.4 | 2.50 | 0.00 |
| Horizontal scanner | - | 4.5 | 2.00 | 4.60 |
| Spherical mirror # 3 | 250 | 50.8 | 0.00 | 3.30 |
| Spherical mirror # 4 | 500 | 50.8 | 2.00 | 0.00 |
| Vertical scanner | - | 9.5 | 3.00 | -3.30 |
| Spherical mirror # 5 | 500 | 50.8 | 0.00 | -3.30 |
| Spherical mirror # 6 | 500 | 50.8 | 3.30 | 0.00 |
| Deformable mirror | - | 7.2 | 3.30 | 0.00 |
| Spherical mirror # 7 | 500 | 50.8 | 1.00 | 2.50 |
| Spherical mirror # 8 | 500 | 50.8 | 3.00 | 0.00 |

**Fig. 1.**
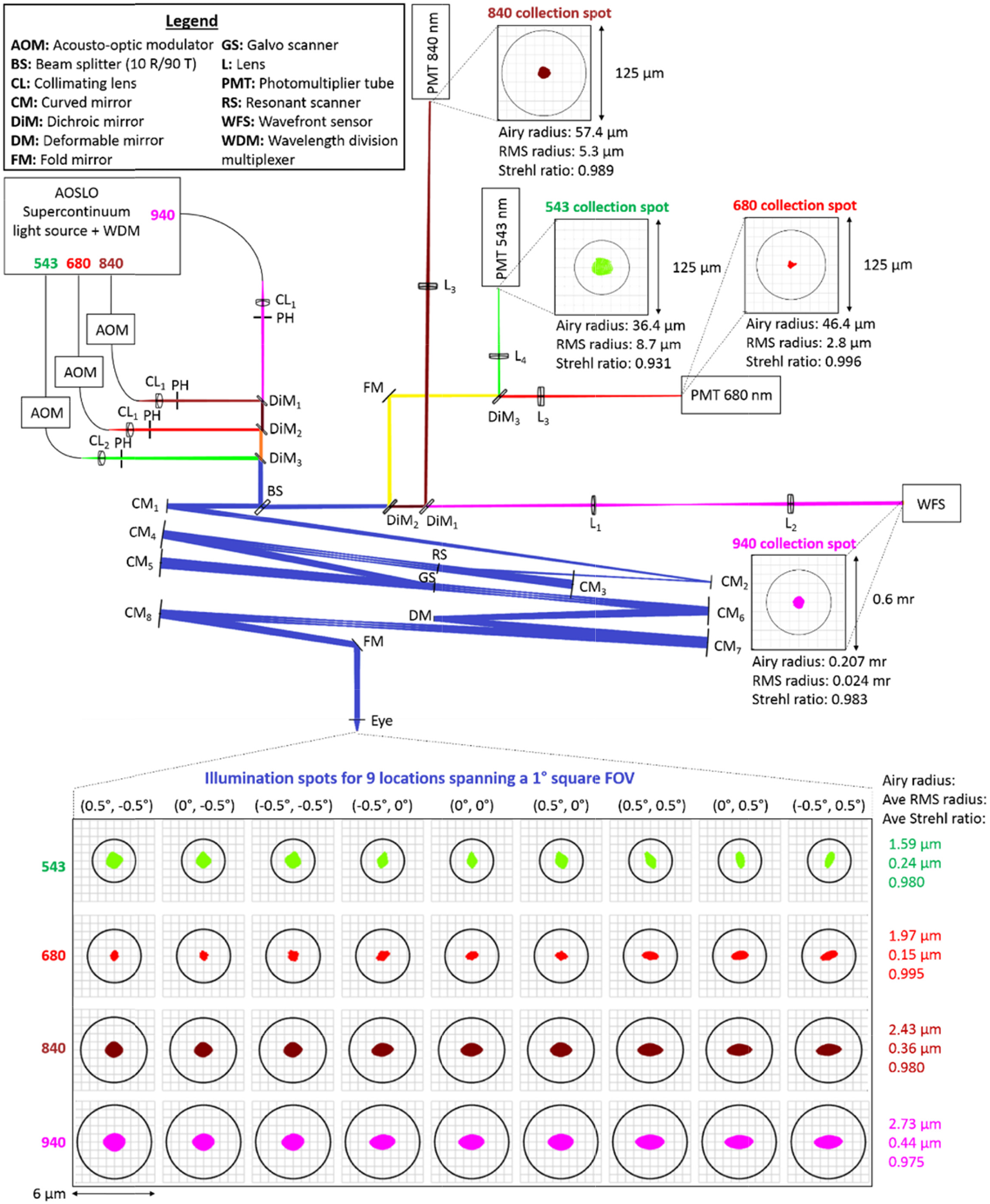
Multi-spectral AOSLO optical design and schematic. All optical components are labeled and described in the legend. Green, red, maroon, and magenta paths correspond to the optical paths of the 54 nm, 680 nm, 840 nm, and 940 nm channels, respectively. Paths with combined channels are blue (all wavelengths), orange (680nm+840nm+940nm), brown (840nm+940nm), and yellow (543nm+680nm). Spot diagrams for all illumination spots spanning a 1° field of view (FOV) are shown below the schematic while those for specific collection channels are placed adjacent to the corresponding collection PMT or WFS as insets. Bene th each spot diagram are the Airy radius, root-mean-square (RMS) spot radius, and Strehl ratio. The model eye consists of a paraxial lens with a 16.7 mm focal length in air.

To minimize aberrations in the collection path, we optimized the optics following the wedge plate beamsplitter in transmission (including the wedge plate beamsplitter itself) and fixed all optical components that had an effect on the illumination path. From our initial analysis, we determined that the wedge plate beamsplitter had the largest contribution to the collection path aberrations. To minimize aberrations induced by this beamsplitter, we optimized for the wedge angle and incident beam size (within optomechanical constraints and beamsplitter availability) to produce a minimum spot size across all spectral channels while allowing a sufficient wedge angle to reject the ghost artifact from the back surface of the beamsplitter (see analysis in Figures 2-4). Based on the simulation results, we chose to use a 0.5° wedge plate beamsplitter (BSX10, Thorlabs Inc, Newton, NJ) positioned along the optical path such that the incident illumination/collection beam size for all spectral channels was minimized given optomechanical constraints.

**Fig. 2.**
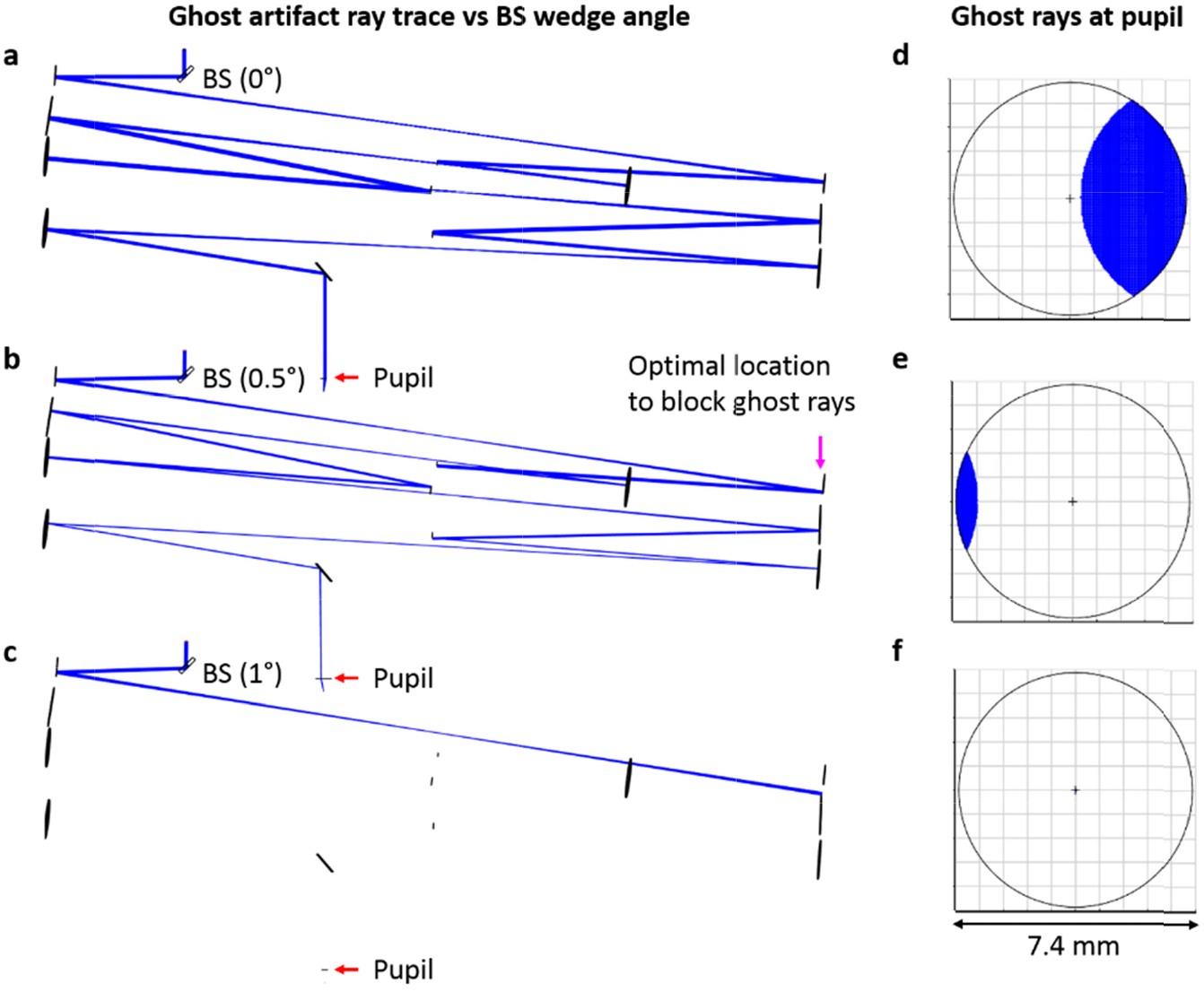
Ghost reflection ray trace analysis for wedge plate beam splitters (BS) of different wedge angles. Ray traces for the light reflecting from the back surface of the wedge beam splitter are shown for wedge angles of 0°, 0.5°, and 1° for (a-c), respectively. The corresponding footprint diagrams for beam profiles at the pupil plane of a model eye are given in (d-f), demonstrating decreasing amounts of the ghost reflection with increases in wedge angle. The magenta arrow in b) indicates a location in the non-scanning portion of the AOSLO system for which an iris can be placed to completely block the ghost artifact without affecting the imaging optical pat of the AOSLO.

**Fig. 3.**
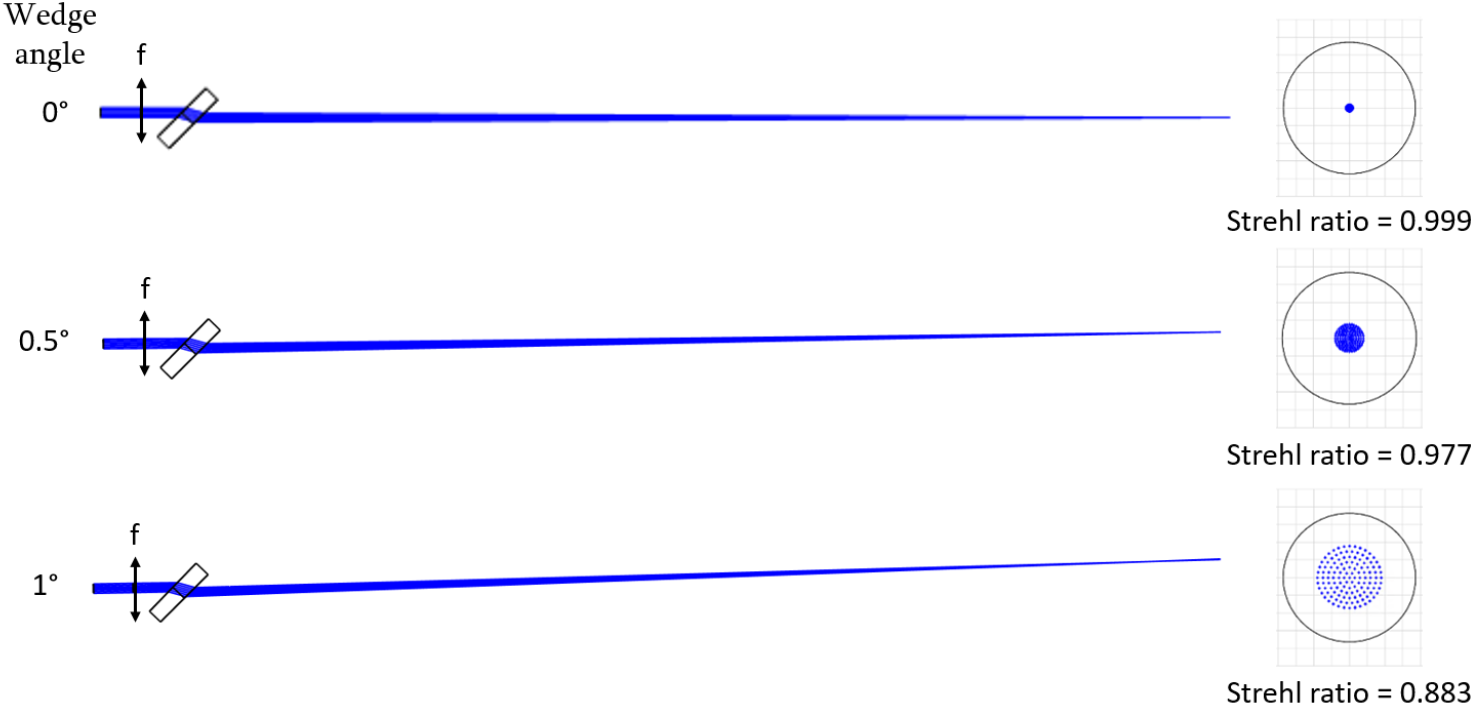
Wedge angle’s effect on aberrations of a converging beam. Schematics on the left each start with a 3.6 mm diameter incident beam of 550 nm light that transmits through a paraxial lens with f = 400 mm (corresponding to a vergence of 0.625 D if preceded by a transverse magnification of ½). Light after the lens focuses through 6 mm thick wedge plate beam splitter made of fused silica and tilted at 45°. The wedge angle is varied from 0° to 1° (from top row to bottom row) and the corresponding spot diagrams at the focal plane are shown at the right.

**Fig. 4.**
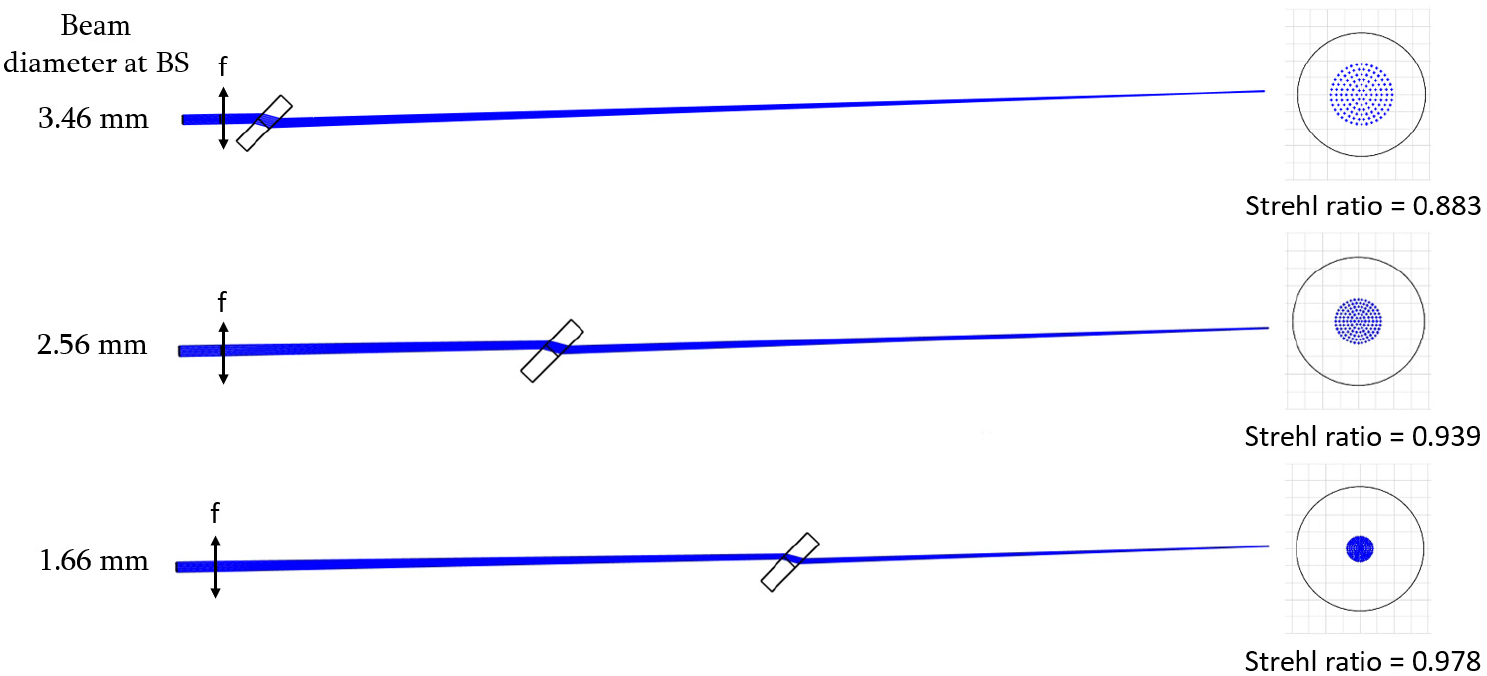
Beamsplitter position’s effect on aberrations of a converging beam. Simulation setups are identical to those in Fig. 3 except that the wedge angle of the beamsplitter is fixed at 1° and the beamsplitter position is altered instead. The schematic shown in the second row has a beamsplitter placed 100 mm closer to the focal plane than that of the schematic in the first row while the beamsplitter in the schematic of the third row is placed 200 mm closer. The beam diameter at the plane of the beam splitter is given to the left of the schematics while the corresponding spot diagrams are shown at the right.

After minimizing the aberrations caused by the wedge plate beamsplitter, the resolution of the AOSLO’s collection path was shown to be diffraction-limited (Strehl ratios > 0.8) for all imaging channels and for the WFS channel (see spot diagrams of collection path in Figure 1). To further improve resolution via confocal gating, approximately ~0.5 of the Airy disk diameter (ADD), which refers to the diameter of the first dark ring in the Airy diffraction pattern, were placed at retinal conjugates prior to the photomultiplier tubes (PMT) imaging channels. The 840 nm imaging channel was slightly more confocal with a pinhole equating to 0.44 ADD, while the 680 nm channel and the 543 nm channel were both collecting 0.54 ADD. For imaging channels with low SNR (e.g. the 543 nm imaging channel), a longer acquisition period was used to allow for robust image registration. To visualize the aberrations at the pupil planes in the system, and therefore at the WFS, we show footprint diagrams illustrating the real ray coordinates throughout the pupil for all scan configurations of the AOSLO (see Figure 5). Here the simulation shows that the deviation in real ray coordinates for all scan positions at the eye and WFS in this AOSLO design (even in the worst case) lie well within the extent of a single lenslet indicating that the wavefront measurements are sampling-limited rather than aberration-limited.

**Fig. 5.**
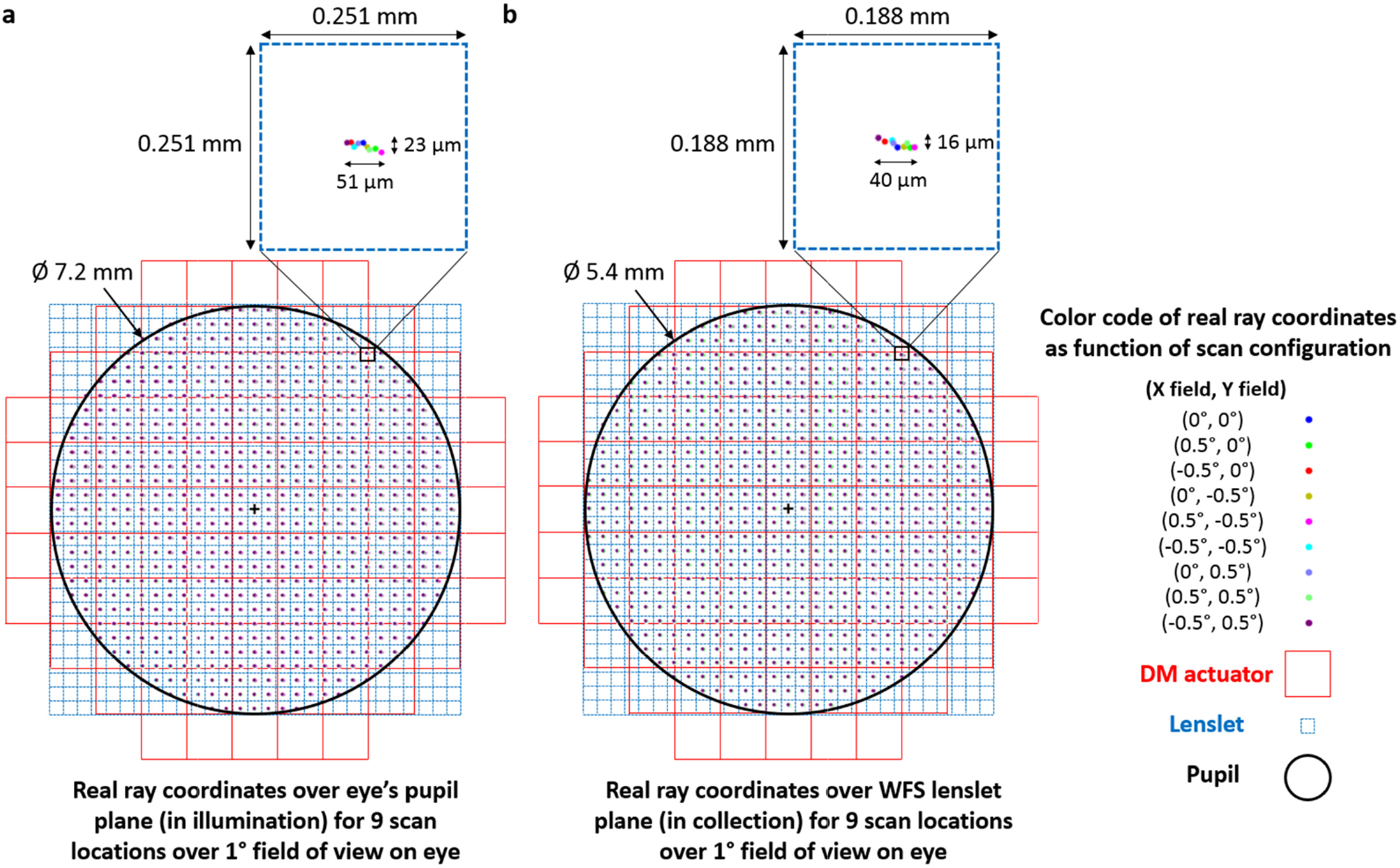
Footprint diagrams show the real ray coordinates over the eye’s pupil (a) and WFS lenslet plane (b) for 9 scan locations spanning a 1° square FOV on the eye. A magnification-corrected image of the system’s pupil, lenslet array, and DM actuator array are overlaid on top of the footprint diagrams revealing that the wavefront measurement is mainly limited by the sampling density dictated by the WFS’s lenslet array rather than the combination of other effects such as pupil aberrations, distortion, and wobble in the illumination and collection paths. The sampling density of the real ray coordinates for the footprint diagrams was chosen to roughly correspond to the magnification-corrected lenslet spacing at the conjugate planes shown in a-b). Magnified insets of a lenslet to the top right of the pupil (where the optical aberrations are worst) are shown at the top right of (a) and (b).

### 2.2 Mechanical design and system construction

The optomechanical design of the AOSLO was designed to prevent vignetting of the optical path, avoid overlapping of optomechanics, and facilitate the optical alignment. The optical path was exported from Zemax optical design software as a solid model to a computer aided design software, Solidworks (Dassault Systemes Solidworks Corp, Waltham, MA), in which the optomechanics were assembled to fit the optical path. In order to aid the alignment, the optical path was oriented such that the en ry beam at the beamsplitter and the exiting beam at the eye were both orthonormal to the table. With this implementation, the threaded holes of the optical table could be utilized to perform accurate and repeatable alignment checks. Each optomechanical component was created or imported (if available from the vendor) into the AOSLO solid model assembly. Kinematic stages and mounts within the AOSLO assembly were designed as sub-assemblies that could be adjusted within its mechanical constraints. Accurate modeling of the kinematic stages allowed the AOSLO solid model to be directly referenced for accurate alignment of optomechanics and aided in the selection of kinematic components to ensure they had sufficient degrees of freedom and range of adjustment to support the optical design.

The WFS optomechanics were custom-built to align a lenslet array and camera (specified in Table 1) with sufficient degrees of freedom for external calibration without sacrificing stability. The WFS camera was anchored to a custom-built adapter plate resting on a tip-tilt kinematic mount which was used to ensure the camera’s sensor was perpendicular to the lenslet array. The lenslet array was held by a high precision rotation mount and attached to a flexure XY translation stage for small, stable adjustments to align the lenslet array to the desired axes of the camera. The WFS module was integrated into the AOSLO with two post holders mounted on a linear translation stage to facilitate system integration and alignment.

In order to accurately construct and align the multi-spectral AOSLO system, we used the optomechanical design to create a stencil for the placement of each optomechanical component on the optical table. As shown in Figure 6(a-b), the spatial coordinates of each post’s location was exported into an Adobe Illustrator drawing shown in Figure 6c. To assure the correct component placement relative to the optical table, multiple reference holes (corresponding to several threaded holes on the optical table) were added to the drawing. The drawing was then laser cut onto a polycarbonate sheet shown in Figure 6d and applied as a stencil to tra e each comp nent’s location onto the optical table using a pencil depicted in Figure 6e. For posts in the design that could be held at a variable height via post holders, their alignment was determined using virtual calipers from the solid model and implemented with sub-millimeter precision using digital calipers. To reduce ambiguity in the optical path length between optics held by kinematic positioners (especially those that allow for a large amount of piston), the pegs for each kinematic mount in the system were coarsely set to the position dictated by the solid model before the final optical alignment.

**Fig. 6.**
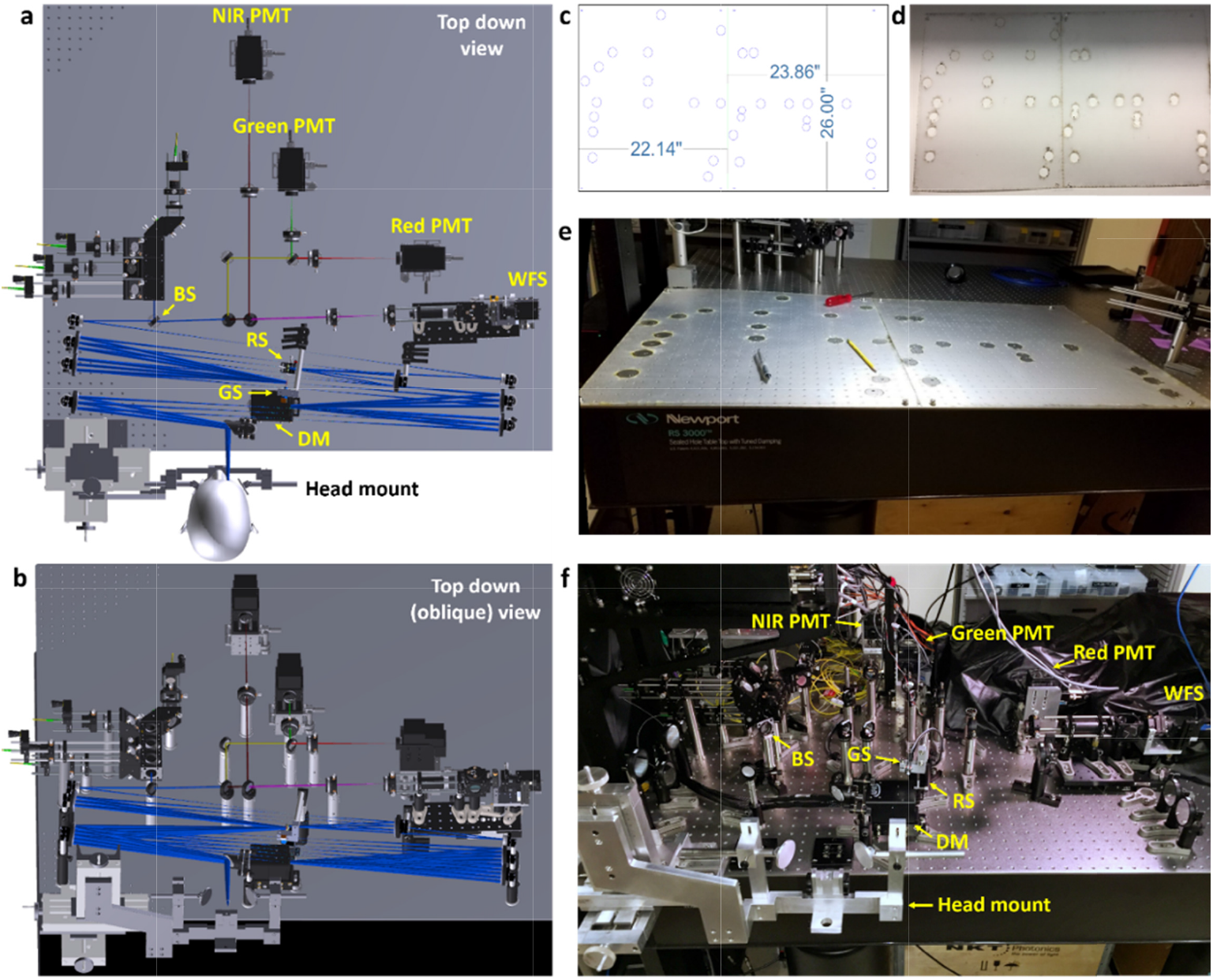
Mechanical design and fabrication of the multi-color AOSLO system. a) Top down view of the optomechanical model of the system within Solidworks. b) Oblique view of the optomechanical model. c) Stencil design indicating the post placement as dictated by the solidmodel shown in a-b). d) Fabricated stencil. e) Stencil applied to optical table. f) Fully-fabricated and aligned multi-color AOSLO system.

### 2.3 Alignment procedure for the AOLSO light delivery and collection

In order to take full advantage of AO correction, the optical path for each imaging channel must be coincident with that of the wavefront sensing spectral channel. Spectral channels were coaligned by making image and pupil planes coincident across all channels. To allow for the alignment of an image plane without misaligning the corresponding pupil plane, we placed the initial collimating lens for each spectral channel on an XY translation stage with a separate iris placed in the front focal plane. With this design, the image plane for any spectral channel could be fine-tuned by shifting the collimator lateral position without affecting pupil position.

To bring all spectral channels into focus at the same plane as the retina, the AOSLO requires optical pre-compensation of the eye’s longitudinal chromatic aberration (LCA). We implemented compensation by adjusting the input vergences of each spectral channel. In this AOSLO design, we chose the 680 nm spectral channel as the reference channel with 0 D vergence making this path collimated prior to the wedge plate beam splitter and at the eye. With 680 nm as a reference, the LCA of the eye was calculated to have a vergence difference of −0.607 D at 543 nm, 0 D at 680 nm, +0.392 D at 840 nm, and +0.5516 D at 940 nm [8,10]. This vergence shift was implemented for each spectral channel by shifting the axial position of the channel’s input fiber such that the image plane for a model eye with a focal length of 100 mm would be 106.46 mm at 543 nm, 100 mm at 680 nm, 96.22 mm at 840 nm, and 94.77 mm at 940 nm. To aid in alignment, a camera was placed at the calculated image plane of the model eye for each spectral channel and the fiber was precisely positioned to minimize the spot size seen on the camera.

### 2.4 Wavefront sensorless adaptive optics to correct static system aberrations

To improve the imaging performance of the AOSLO system and make it robust to minor system aberrations, we utilized wavefront sensorless adaptive optics to determine the optimal wavefront shape that maximizes detected light. Our implementation of sensorless adaptive optics is similar to that described by Hofer *et al.* [21] and applies random perturbations to the deformable mirror shape to optimize for the AOSLO image’s mean intensity rather than the wavefront sensor readings. By using a small confocal pinhole (with a sub-Airy disk diameter), aberrated and out-of-focus light is rejected by the confocal pinhole and the wavefront shape that optimizes for mean intensity also corresponds to the wavefront with minimal aberrations. Once the deformable mirror shape is optimized based on the detected image’s mean intensity, the residual wavefront reading on the WFS is recorded and is assigned as the target shape for subsequent WFS-based AO correction.

### 2.5 Image processing and human subject protocol

The system components and hardware customized for this system are listed in Table 1. The custom image acquisition software and AO software are same as that of our previous AOSLO system and have been previously described in [22, 23]. The image registration techniques used for stabilizing eye motion was a strip-based cross correlation method previously described in [24, 25].

The University of California Berkeley Institutional Review Board approved this research, and subjects signed an informed consent form before participation. All experimental procedures adhered to the tenets of the Declaration of Helsinki. Mydriasis and cycloplegia were achieved with 1% tropicamide and 2.5% phenylephrine ophthalmic solutions before each experimental session. Subjects bit into a dental impression mount affixed to an XYZ translation stage to align and stabilize the eye and head position. Both subjects (20112L and 20076R) were healthy young adult volunteers. Structural imaging was performed on both subjects using the 543 nm, 680 nm, and 840 nm imaging channels and the 940 nm WFS channel for AO correction.

## 3. Results

After constructing and aligning the multi-spectral AOSLO system, the optical resolution for each spectral channel was quantified using a phase retrieval technique and a through-focus stack of intensity images [26, 27]. The phase retrieval algorithm utilized through-focus intensity images to iteratively solve for the complex PSF using the Gerchberg-saxton algorithm [28]. A Fourier transform of the complex PSF was then used to reconstruct the wavefront and determine optical quality metrics like Strehl Ratio, which we used to evaluate resolution for each spectral channel (see Table 3).

**Table 3.** Strehl Ratio measurements for the AOSLO’s illumination and collection paths per spectral channel.

| Spectral channel | Illumination | Collection |
| --- | --- | --- |
| 543 nm | 0.95 | 0.94 |
| 680 nm | 0.93 | 0.99 |
| 840 nm | 0.99 | 0.96 |
| 940 nm | 0.97 | 0.98 |

The optical design and system alignment were further validated after measuring the system’s collection efficiency per spectral channel as a function of confocal pinhole size (see Figure 7). In this experiment, the sample was a model eye with paper as the retina and the illumination profile at the pupil plane was a top hat with a circular aperture. In the absence of scattering effects, the spatial profile of the light incident on the confocal pinhole is expected to follow that of the double-pass point spread function’s encircled energy [29], which follows the black curve on Figure 7. However, scattering within paper is expected to further broaden the light distribution at the confocal pinhole with its own point spread function, which has been shown to approximately follow a Lorentzian function [30] (see dashed lines in Figure 7). To measure the paper’s scattering point spread function, a separate simplified setup with the same illumination profile and model eye from the AOSLO was used to measure the collection efficiency per imaging channel vs confocal pinhole size (see dense dotted lines in Figure 7). Comparing the encircled energy for the multi-spectral AOSLO with the measured encircled energy of the simplified setup, we see that our results match quite closely indicating the multi-spectral AOSLO’s near-theoretical collection efficiency across all imaging channels. As reference, we also plot related work from Sredar *et al.* [31] and an encircled energy curve resulting from the convolution between the theoretical double-pass point spread function (without scattering) and a 2 μm FWHM Lorentzian for each imaging channel. This latter plot matches closely to the encircled energy measurements of our simplified setup indicating that the paper of our model eye has a PSF that approximately equals to a 2 μm FWHM Lorentzian function.

**Fig. 7.**
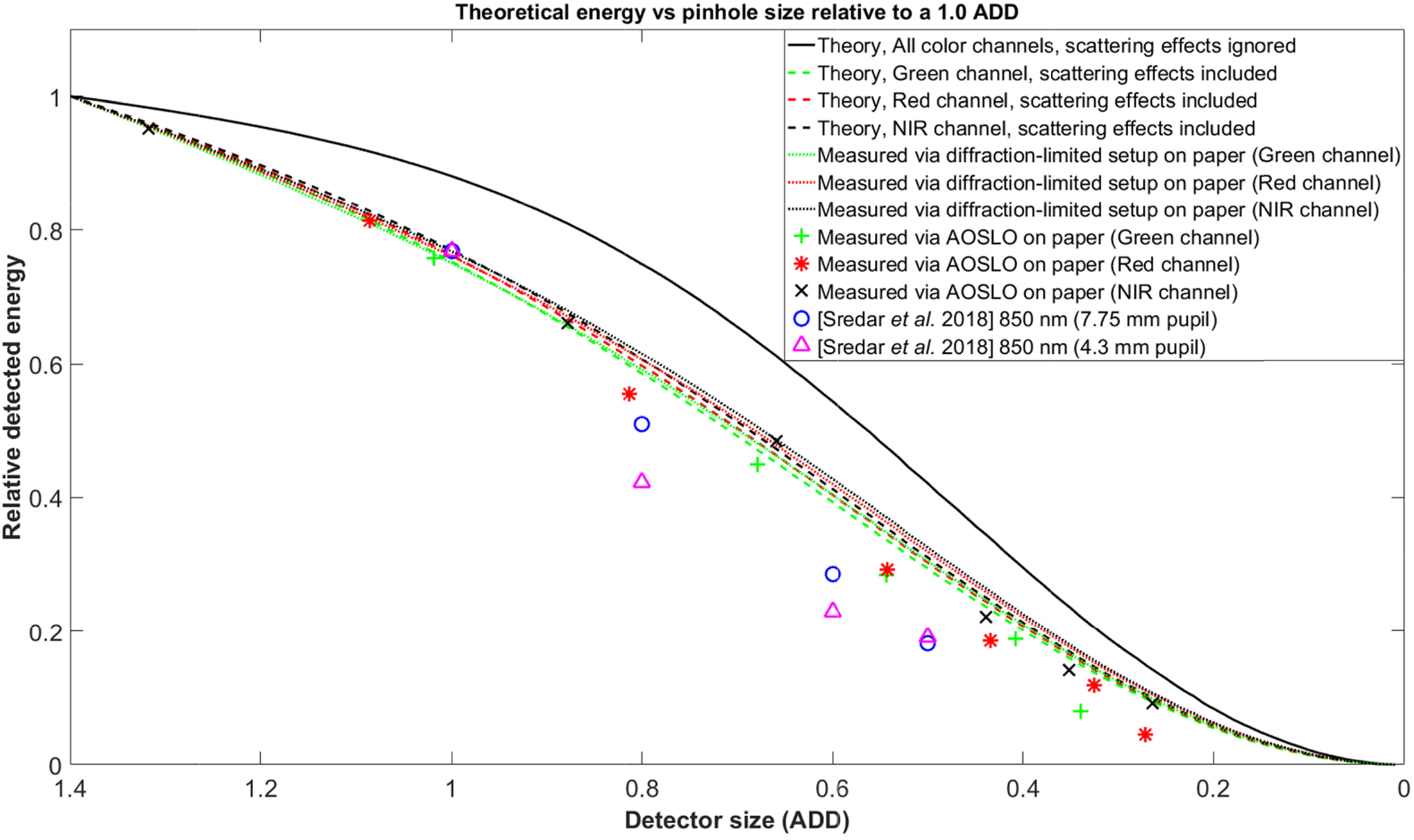
Theoretical and experimental energy at the confocal pinhole for the AOSLO and a separate diffraction-limited setup when imaging the same model eye and spectral imaging channels. Theoretical estimates without scattering were calculated by integrating the double-pass point spread function at the plane of the confocal pinhole for different pinhole sizes. Theoretical estimates incorporating scattering were calculated by convolving the double-pass point spread functions by a 2 μm FWHM Lorentzian function and integrating the result for different pinhole sizes. All curves were normalized to unity for a confocal pinhole size of 1.4 ADD The detected energy of the largest pinhole size for each spectral channel was normalized to the value of the curve generated from measurements using the diffraction-limited setup on paper for the corresponding spectral channel and pinhole size.

To test the performance of our multi-spectral AOSLO system for imaging the human eye, we obtained preliminary imaging results from two emmetropic, healthy human adult volunteers at the fovea after artificial dilation. Videos of the foveal cone mosaic were acquired in all imaging channels at 30 Hz with a 0.83° FOV and averaged for 10 seconds to produce the averaged images shown in Figure 8. The illumination powers at the eye and ratio of the maximum permissible limit for a 0.75° FOV were 4.6 μW (0.42), 69 μW (0.16), 87 μW (0.08), and 94 μW (0.05) for the 543 nm, 680 nm, 840 nm, and 940 nm spectral channels, respectively. For all imaging sessions, only one imaging channel was utilized at a time together with the wavefront sensing channel, although simultaneous imaging with all channels could be permitted under the maximum permissible safety limits [32, 33].

**Fig. 8.**
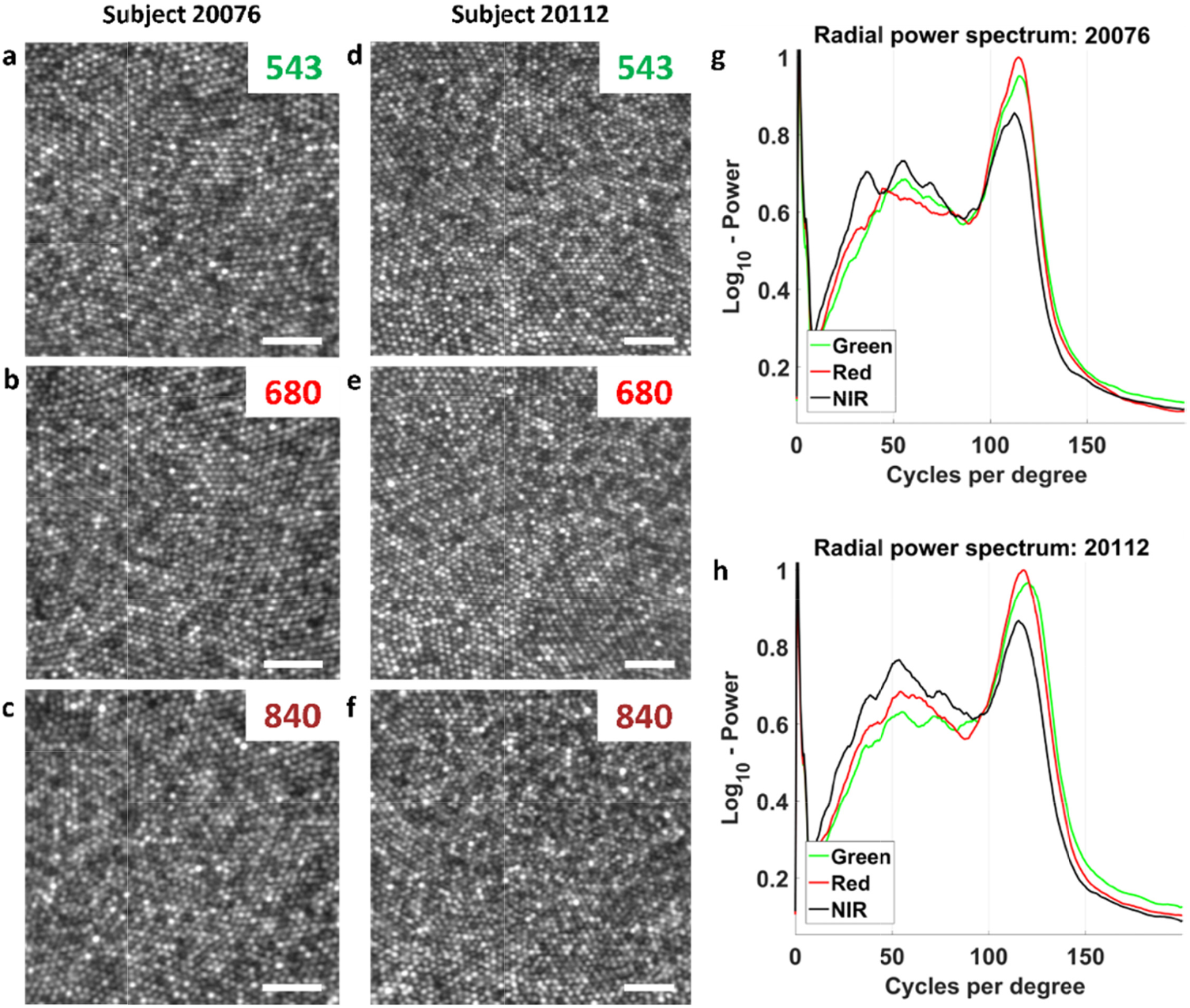
Foveal imaging results across all spectral channels from healthy hyperopic (a-c, Subject 20076) and emmetropic (d-f, Subject 20112) volunteers after artificial dilation and ~0.5 ADD pinhole configuration. a,d) 543 nm channel image. b,e) 680 nm channel image. c,f) 840 nm channel image. g,h) Radial power spectrums across all imaging channels for Subjects’ 20076 and 20112, respectively. Scale bars, 20 μm.

For the spectral analysis, 100 frames were registered for motion using strip-wise cross correlation algorithm and averaged for the 680 nm and 840 nm spectral channels. To account for the low power used in the 543 nm imaging channel, a total of 300 frames were registered and averaged for the spectral analysis. The images were all cropped to the same region of interest and t e histograms of the images were normalized across the full bit depth to replicate the same contrast.

The spatial frequency information from images of different spectral channels was quantitatively compared for both subjects using the radially-integrated power spectrum (Figure 8g, 8h). The power spectrum from each channel was normalized to the area within each curve such that the same amount of information is encapsulated in each distribution. As expected by diffraction-theory [34], the 543 and 680 nm imaging channels demonstrated sharper images with higher resolution and their radial power spectrums exhibited a larger peak at the cone-packing density (~110-125 Cycles/degree) than that of the 840 nm imaging channel.

However, the benefits for the shortest wavelength, 543 nm, was not fully realized compared to 680 nm with a 0.5 ADD pinhole configuration. There are several possible reasons for this. The primary cause, put forth in the introduction, was that the different optical paths between 940 and 543 nm light might have resulted in just enough high order aberration differences to preclude an optimal correction. But other factors might also contribute: The lower power that was used for green light imaging might have resulted in slightly poorer eye motion correction and consequent image registration. Finally, the high PMT detector gain required for green light detection might have resulted in more high frequency noise (~150-200 cycles/degree) compared to the other channel. An increase in power in this part of the spectrum would affect the power spectrum normalization in a way to reduce the relative height of its peak. Nevertheless, the images remain very good at short wavelengths despite all possible reasons for it not to be.

The practical implications are that one can achieve foveal cone resolution in both 543 nm and 680 nm spectral channel when the correction is informed by the measurements of the eye’s ocular aberrations at 940 nm. This study shows the practical limits of the different spectral channels and that both 680 nm and 543 nm imaging can be used for structural foveal imaging, psychophysical experiments in the fovea [11] and measuring and correcting for TCA [13,14].

## 4. Conclusions

We have demonstrated a multi-spectral AOSLO design with diffraction-limited illumination and collection to achieve high-resolution, high-throughput retinal imaging. After constructing and validating the AOSLO performance, images were acquired at the foveal center from two healthy subjects to demonstrate the system’s capability to visualize foveal photoreceptors in all imaging channels with wavefront correction based on a separate spectral channel. The use of this methodology and system design may provide increased collection efficiencies in other SLO or AOSLO designs that employ large vergence ranges over multiple spectral channels.

